# Using total RNA quality metrics for time since deposition estimates in degrading bloodstains

**DOI:** 10.1101/2021.04.28.441847

**Authors:** Colin I. Elliott, Theresa E. Stotesbury, Aaron B.A. Shafer

## Abstract

Determining the time since deposition (TSD) of bloodstains would provide forensic scientists with critical information regarding the timeline of the events involving bloodshed. The physicochemical changes occurring to biomolecules as a bloodstain degrades can be used to approximate the TSD of bloodstains. Our study aims to quantify the timewise degradation trends and temperature dependence found in total RNA from bloodstains without the use of amplification, expanding the scope of the RNA TSD research which has previously targeted mRNA molecules. Whole bovine blood was stored in plastic microcentrifuge tubes at 21°C or 4°C and tested over different timepoints spanning one week. Nine RNA metrics were visually assessed and quantified using linear and mixed models; the RNA Integrity Number equivalent (RINe) and the DV200 demonstrated strong negative trends over time and statistical independence. The RINe model fit was high (R^2^ = 0.60), and while including the biological replicate as a random effect increased the fit for all RNA metrics, no significant differences were found between biological replicates stored at the same temperature for the RINe and DV200 metrics. Importantly, this suggests that these standardized metrics can likely be directly compared between scenarios and individuals, with DV200 having an inflection point at ∼28 hrs. This study provides a novel approach for blood TSD estimates, producing metrics that are not affected by inter-individual variation and improving our understanding of the rapid degradation occurring in bloodstains.

**HIGHLIGHTS:**

- Amplification-free analysis of total RNA in degrading bloodstains.
- Short-term RNA degradation assessment using high-resolution size measurements.
- Total RNA quality and quantity metrics were assessed across a one-week.
- Total RNA quality metrics demonstrated the strongest timewise trends.
- Biological replicates produced similar results for RNA quality metrics.

## Introduction

Accurately determining the age of biological evidence, including soft tissue and bodily fluids, remains a complex task in forensic investigations (1,2). Bloodstained evidence is commonly observed at the scenes of crimes involving bloodshed, and determining the age, or time since deposition (TSD) of a bloodstain, would provide context in criminal investigations, including the development of investigative leads and the assessment of alibis (1–4). Currently, bloodstains observed at crime scenes can be subjected to bloodstain pattern analysis, which provides potential interpretations of the physical events that gave rise to the bloodshed (5).

DNA profiling, used for source attribution purposes, can be carried out from collected bloodstains (6–9); however, these tools do not provide TSD estimates of blood, requiring additional analyses. Bloodstains degrade in a series of complex processes, physical and chemical changes are influenced by many factors, starting from the point of their formation on a surface onwards (10–16). Previous studies have developed chemometric (2,13,14,17) and genetic methods (3,18–20) that estimate the TSD of bloodstains in controlled laboratory conditions. While these methods produce time-dependent responses, studies generally report high variability in responses between samples, biological replicates and environmental conditions, suggesting that the accuracy of TSD estimates might be increased by combining metrics obtained from different biomolecules and/or using methods that are insensitive to biological replicate variation (1,3,13,14,17–19,21).

*Ex-vivo*, blood undergoes a series of time-dependent physicochemical and morphological changes, such as hemoglobin oxidation and conformational changes (13,22), drying (12,23), clotting (24,25), and DNA and RNA degradation (3,15,26–28). Total RNA encompasses ribosomal RNAs (rRNA), messenger RNAs (mRNA), transfer RNAs (tRNA) and microRNAs, among others (29,30). Eukaryotic rRNA makes up for 80% of the total RNA in a cell and consists of 5S, 5.8S, 18S and 28S rRNA (31). Messenger RNA expression patterns have been used to identify body fluids found at crime scenes (32,33) and might also correlate with time (34). Fu *et al.* (20) found that, when analyzing blood-specific mRNA transcripts, TSD for bloodstains could be estimated to within a period of 2 to 6 weeks, depending on the TSD of the sample. Anderson *et al.* (3) distinguished 6-day old blood from fresh and aged (30 days) blood, while Lech *et al.* (18) classified deposition by three time-of-day periods with high prediction accuracy. High throughput sequencing (HTS) by Weinbrecht *et al.* (19) demonstrated that mRNA transcripts decreased over time, with blood-specific transcripts detected up to 12 months after deposition. Salzmann *et al.* (35) also used HTS to investigate the timewise changes in the quality of the amplified RNA; the authors observed a marked decrease in mRNA transcript integrity six months after deposition, highlighting the potential applicability of total RNA quality metrics for TSD purposes (35). Collectively, these studies demonstrated the timewise degradation of small RNA species in bloodstains and highlighted the surprising long-term survival of RNA, supporting the applicability and implementation of techniques targeting nucleic acid degradation in time series. We ask if features of quick RNA degradation can be captured in the same way.

Most RNA studies to date have used PCR amplification to target specific mRNAs and how they change in a degrading bloodstain over a long period of time (3,19,20,35). The vast majority of mRNA molecules have a half-life of just minutes (36); therefore, genomic DNA contamination is a concern in experiments where amplification is used to target specific Mrna (37). Amplification-free analysis of RNA might provide insight into the naturally occurring RNA degradation in blood without the risk of introducing any amplification biases or stochastic effects seen while using RT-PCR and RT-qPCR (36,38,39). Accordingly, our study takes a holistic, non-targeted and amplification-free approach by analyzing the quality (ribosomal ratios, RNA integrity number equivalent; RINe, DV200) and quantity metrics (RNA size ranges, total concentration, rRNA peak concentration) of total RNA in bloodstains, while simultaneously quantifying the effect of temperature on RNA degradation and TSD models. Not only does focusing on total RNA have the advantage of eliminating the stochastic effects and amplification biases caused by reverse transcriptase and PCR, it can also circumvent potential PCR inhibition from the heme in blood (38–40).

## Materials and Methods

### Blood sample collection

Bovine blood was collected from Otonabee Meat Packers, an abattoir in Peterborough, Ontario, Canada. Acid dextrose anticoagulant solution A (ACD-A) was added to bovine blood at a concentration of 12.5% v/v (41). Blood treated with ACD anticoagulants has demonstrated better nucleic acid extraction efficiencies than heparin-based anticoagulants and has not been shown to affect the cell counts of the different blood components (42). The ACD-A used in this study was not shown to impact RNA metrics such as concentration and quality over time (Table S1) and allowed for more accurate pipetting of the blood. ACD-A was made by dissolving 6.6 grams of sodium citric dextrose, 2.4 grams of citric acid, and 6.68 grams of dextrose anhydrous, all purchased from ACP Chemicals Inc., in 500 mL of distilled water. Bovine blood was collected in an amber Nalgene bottle containing the anticoagulant, which was subsequently placed on ice for transportation back to the laboratory (approximately 15 minutes).

### Blood deposition

50 μL of blood was deposited into 1.5 mL plastic microcentrifuge tubes. Samples were left to air dry by keeping the microcentrifuge tubes lids open until RNA extraction. Samples were stored at one of two temperatures: room temperature (∼ 21°C) or refrigerated at 4°C for a designated amount of time. The room temperature storage condition had an approximate 40% relative humidity (RH), while the RH was likely slightly greater for samples stored at 4°C. Five time-series experiments were performed, three at 21°C and two at 4°C; blood from a different cow (biological replicate) was used for each experiment. RNA extractions were completed at 0 hours (time of initial pipetting; ∼ 30 minutes post-blood collection) and across various timepoints (Table 1); the goal here was to compare the timewise degradation trends for blood deposited at different temperatures. Five technical replicates were used for each of the timepoints, resulting in a total of 200 assayed samples.

**Table 1.** Overview of the time series conducted and each experiment’s storage temperature for the bloodstains. Each biological replicate represents a different cow.

| Biological Replicate | Temperature | Sample Size | Timepoints (hours after deposition) |
| --- | --- | --- | --- |
| A | 21°C | 50 | 0,1.5,3,6,9,24,48,72,96,168 |
| B | 21°C | 50 | 0,1.5,3,6,9,24,36,48,72,96 |
| C | 21°C | 30 | 0,6,20,24,28,36 |
| D | 4°C | 35 | 0,3,6,9,24,48,72 |
| E | 4°C | 35 | 0,3,9,24,48,72,96 |

### RNA extraction

Total RNA was extracted from the deposited blood using the PureLink Total RNA Blood Purification kit by Invitrogen (43). The volume of lysis buffer (L5) recommended in the manufacturer’s protocol was doubled to minimize blood clotting. An optional on-column DNase incubation was completed following the manufacturer’s protocol. Total RNA was eluted with 30 μL of RNase-free water and prepared for analysis on the Agilent Technologies 4200 TapeStation (44). 1 μL of the high sensitivity RNA screen tape sample buffer was mixed with 2 μL of the total RNA extracts following the manufacturer’s protocol for the *High Sensitivity RNA ScreenTape Assay* (44).

### Automated gel electrophoresis

RNA was analyzed using high-sensitivity automated gel electrophoresis with the Agilent Technologies 4200 TapeStation (44). The TapeStation is a highly sensitive instrument that detects RNA down to 100 pg/μl while offering a quantitative range beginning at 500 pg/μl (45). It separates different-sized fragments that are tagged with an intercalating fluorescent dye by automated electrophoresis, allowing for the quantification of specifically-sized RNA (45,46). A ladder containing fragments of known size was also loaded next to the samples; the software compared band locations and assigned 18S and 28S rRNA peaks to the RNA samples (47).

### RNA metrics

A series of RNA metrics were collected and examined using the TapeStation Analysis Software A.02.02 (Table S2). Of note, the RINe metric was generated by an algorithm that calculates the ratio of the height of the 18S rRNA peak (in fluorescent units; FU) to the background RNA signal (FU), which represents the signal for fragments found in the fast zone ranging from 200 bp to 1.8 kb (45,47). The RINe metric is a dimensionless number ranging from 1 to 10, with 10 representing intact and high-quality RNA, while a value of 1 represents low-quality and highly fragmented RNA (45,48). Another RNA quality metric, the DV200, represents the percentage of RNA fragments greater than 200 nucleotides (49). A custom RNA bin was made to estimate concentration from 200 bp to 500 bp fragments, presumably representing the shorter mRNAs extracted from our samples (50).

### Statistical analyses

Correlation matrices were created prior to building models to ensure that the response variables (RNA metrics) included in the analyses were not highly correlated to each other (Pearson’s r < |0.70|). Variables showing independence and a correlation to time were used to build a series of linear mixed-effects models, with time (natural log-transformed) and temperature as fixed effects and biological replicate (individual cow) as a random effect. An interaction term between log-transformed time and the temperature was also included in our models. We assigned timepoint zero a value of 0.1 (6 minutes) prior to the logarithm transformation. Effect size, R^2^, and p-values were recorded for each model.

We ran an unpaired two-sample Wilcoxon test, a non-parametric test, to determine whether there was a difference in the biological replicates undergoing the same experimental conditions across all timepoints (i.e. test for between-individual variance in RNA metrics). The non-linearity of the RNA metrics over time was also explored by fitting non-linear curves and assessing the impact on the residual standard error. We note the regressions do not require shared time points between biological replicates to estimate slope, intercept, and inflection points. All statistical analyses and data visualizations were created using R Version 4.0.3. Raw data and scripts are publicly available and deposited at https://gitlab.com/WiDGeT_TrentU.

## Results

We collected nine RNA metrics on 200 deposited bloodstains; four RNA metrics were retained for statistical analyses as they were not correlated to each other (Pearson’s r < |0.70|; Fig. 1). The retained metrics included the RINe, the DV200, 28S/18S rRNA peak area ratio, and total RNA concentration (Table 2).

**Fig. 1-.**
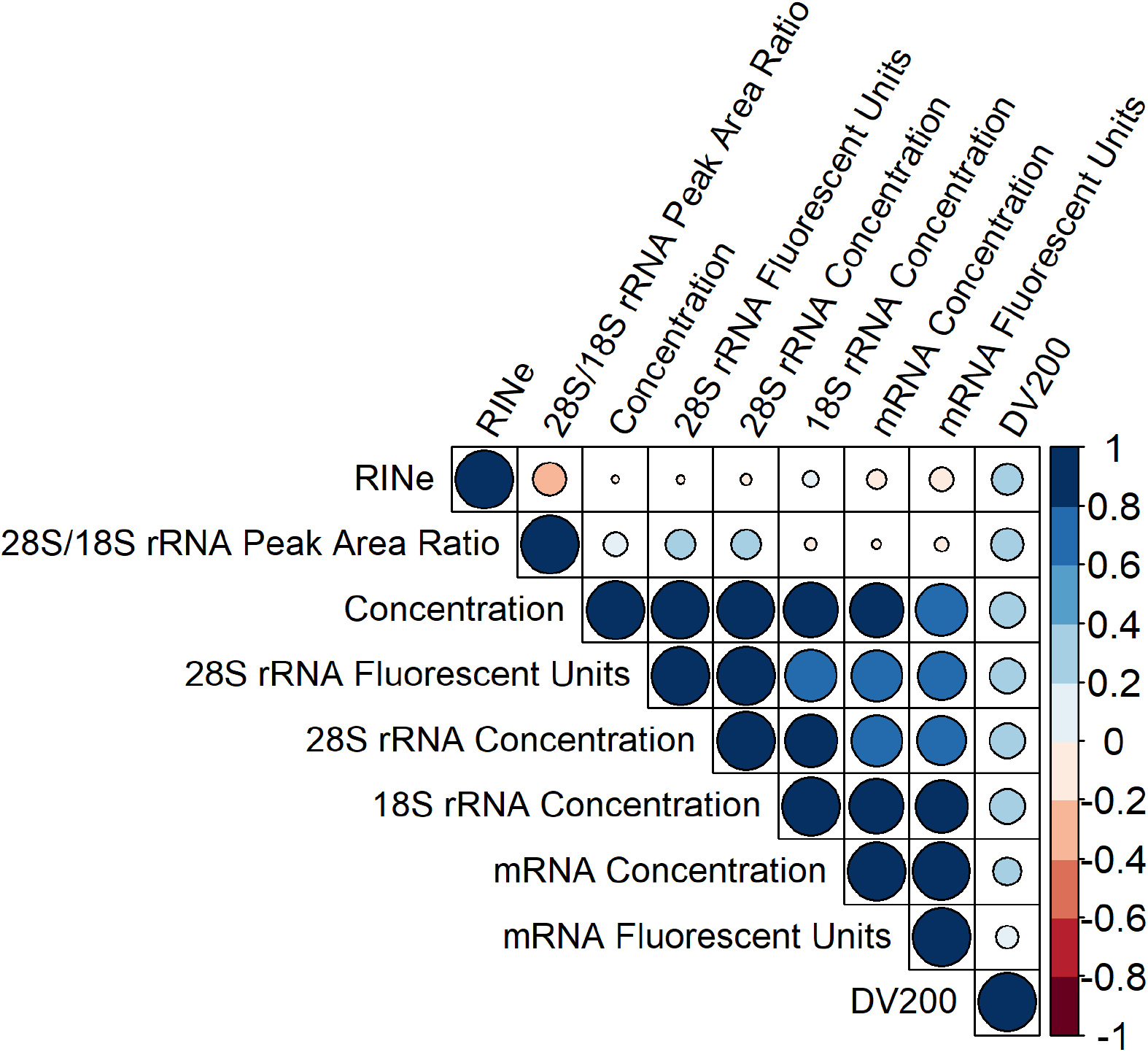
Correlations between the nine RNA metrics obtained from the 4200 TapeStation RNA analysis. The size of the circle increases with a greater correlation between variables. Blue-coloured circles indicate a positive correlation between RNA metrics, while red-coloured circles indicate a negative correlation between them. The shading of the circle increases with a greater correlation, with light colours indicating a weak correlation and darker colours indicating a strong correlation.

**Table 2.** Statistics for the mixed models of the four quantified RNA metrics. Note that there was an interaction between log-transformed time and temperature. Biological replicate was used as a random effect. Each regression was computed individually for each RNA metric and each row represents a separate model. Because of the interaction term, log(Time) in the table represents the relationship between the response variable and time at 21°C; Temperature (4°C) represents the difference between the intercept at both temperatures; and log(Time)*Temperature reflects the difference in correlation between retained RNA metrics and time between 21°C and 4°C. P-values below 0.05 are bolded.

| RNA metrics | Marginal R<sup>2</sup> | Conditional R<sup>2</sup> | log(Time) |  |  | Temperature (4°C) |  |  | log(Time)*Temperature |  |  |
| --- | --- | --- | --- | --- | --- | --- | --- | --- | --- | --- | --- |
|  |  |  | Estimate | Confidence Interval | p-value | Estimate | Confidence Interval | p-value | Estimate | Confidence Interval | p-value |
| RINe | 0.51 | 0.60 | -0.51 | -0.58 – -0.43 | **<0.001** | 0.12 | -0.74 – 0.98 | 0.78 | 0.17 | 0.04-0.29 | **0.01** |
| DV200 | 0.31 | NA | -2.4 | -2.99 – -1.80 | **<0.001** | -0.47 | -3.59 – 2.64 | 0.77 | 1.99 | 1.03 – 2.96 | **<0.001** |
| 28S/18S Peak Area Ratio | 0.061 | 0.24 | 0.34 | -0.21 – 0.89 | 0.22 | 0.35 | -3.50 – 4.20 | 0.86 | 0.19 | -0.54 – 0.91 | 0.62 |
| Concentration | 0.069 | 0.30 | -885.42 | -1305.24 – -465.60 | **<0.001** | -2142.06 | -7554.02 – 3269.90 | 0.44 | 789.89 | 116.22 – 1481.56 | **0.022** |

### RNA Integrity Number equivalent

The RINe metric demonstrated the best fit model (R^2^ = 0.60; Table 2) and a relationship with log-transformed time (Table 2). RINe values were influenced by storage temperature, however, the effect of temperature on RINe varied according to time, as seen by the significant interaction effect in Table 2. The addition of biological replicate (donor) as a random effect slightly increased the model’s goodness-of-fit (marginal vs conditional R^2^ in Table 2); this was driven by a temperature-donor effect, as RINe values obtained from the different biological replicates at the same temperature were not different from each other (Table 3), demonstrating consistency across individuals in the same treatment. The timewise changes in RINe at different temperatures (Fig. 2) showed 21°C caused more degradation (steeper slope) than the samples stored at 4°C (Fig. 2).

**Table 3.**
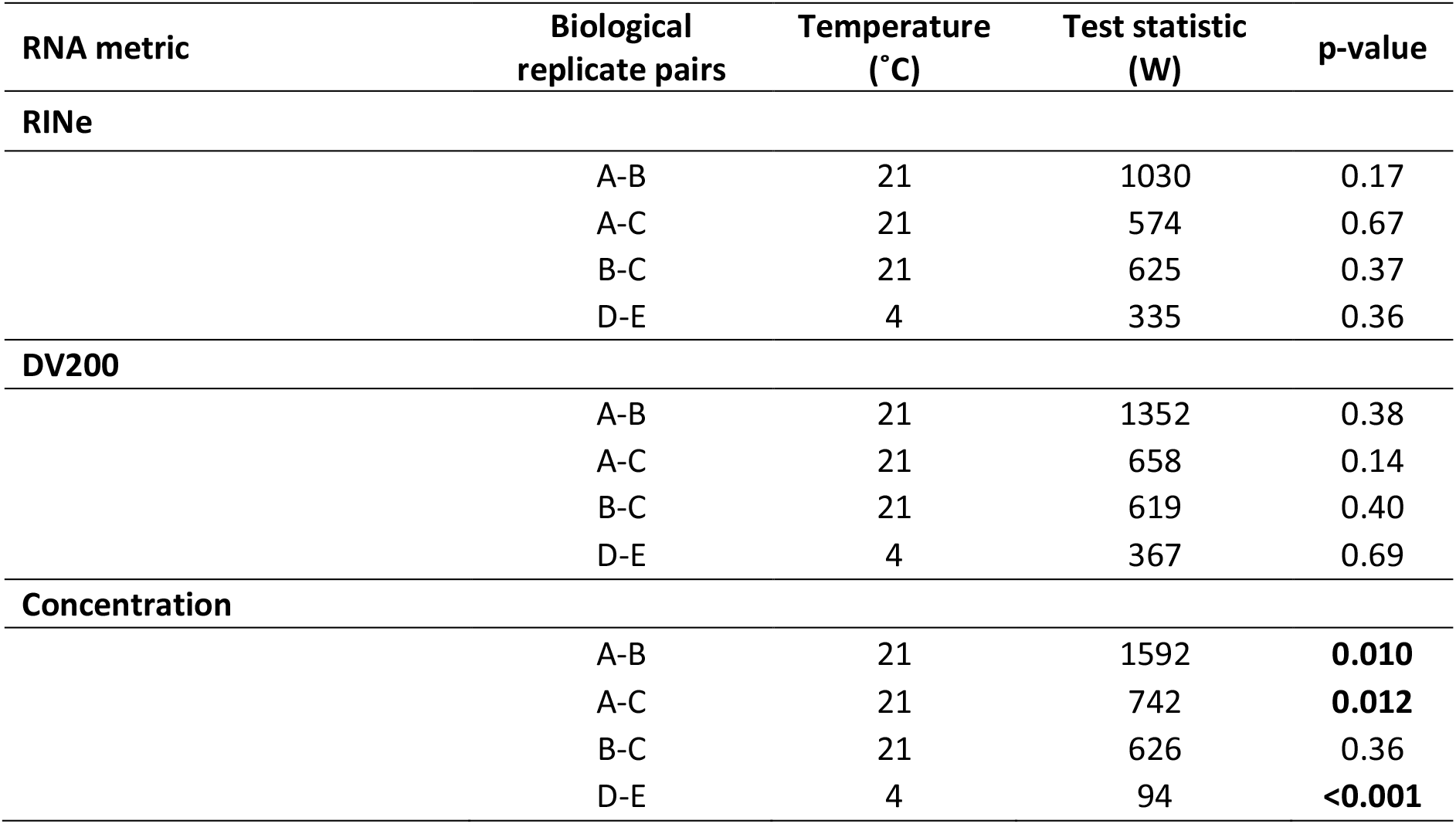
Unpaired two-sample Wilcoxon tests between the different biological replicates for the RINe, DV200 and total RNA concentration. Significant p-values are bolded.

| RNA metric | Biological replicate pairs | Temperature (°C) | Test statistic (W) | p-value |
| --- | --- | --- | --- | --- |
| **RINe** |  |  |  |  |
|  | A-B | 21 | 1030 | 0.17 |
|  | A-C | 21 | 574 | 0.67 |
|  | B-C | 21 | 625 | 0.37 |
|  | D-E | 4 | 335 | 0.36 |
| **DV200** |  |  |  |  |
|  | A-B | 21 | 1352 | 0.38 |
|  | A-C | 21 | 658 | 0.14 |
|  | B-C | 21 | 619 | 0.40 |
|  | D-E | 4 | 367 | 0.69 |
| **Concentration** |  |  |  |  |
|  | A-B | 21 | 1592 | **0.010** |
|  | A-C | 21 | 742 | **0.012** |
|  | B-C | 21 | 626 | 0.36 |
|  | D-E | 4 | 94 | **<0.001** |

**Fig. 2-.**
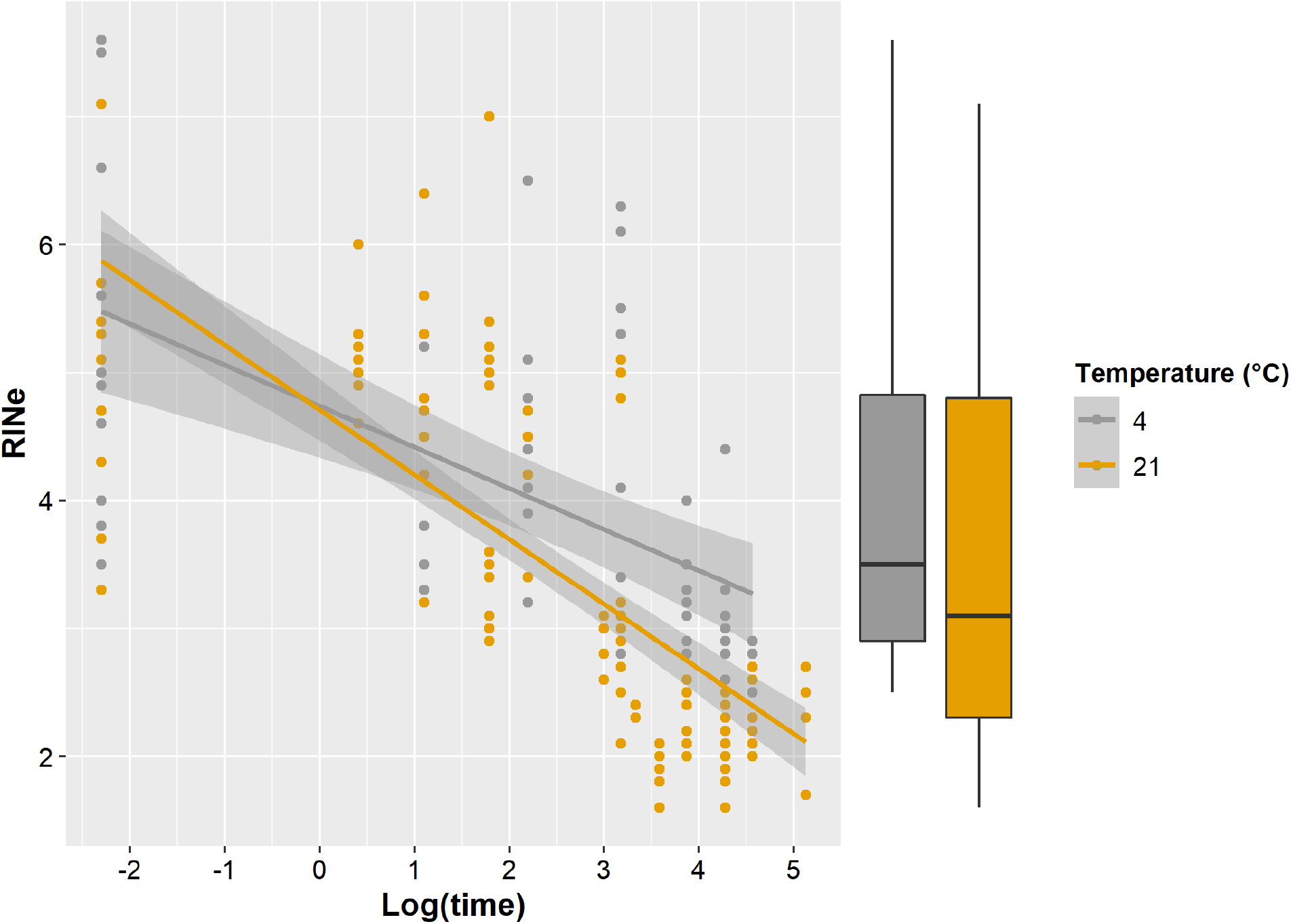
Relationship between RINe and log-transformed time (hours). RINe values are grouped by storage temperature for comparison purposes; points and regression lines are coloured according to storage temperature. Shaded areas around the regression lines represent the standard error. The interaction effect between log-transformed time and temperature is not shown as data is already separated by storage temperature.

### DV200

The DV200 demonstrated the second-best model fit (R^2^ = 0.31); the random effect of biological replicate did not improve the model (Table 2). DV200 showed a statistically significant relationship with time and was influenced by storage temperature, the effect of which varied according to time (Table 2). The positive estimate for the interaction effect indicated that over time, bloodstains kept at 4°C provided greater DV200 values relative to those stored at 21°C (Table 2, Fig. 3). DV200 values obtained from the different biological replicates at the same temperature were not different from each other (Table 3), but there was a difference between DV200 values recovered from bloodstains deposited at 21°C and 4°C (W = 2572, p-value = 0.015; Fig. 4A-B). Bloodstains stored at room temperature exhibited greater levels of fragmentation over time (Fig. S1, Fig. 4A-B). DV200 values from bloodstains deposited at room temperature also exhibited a non-linear trend over time; we determined that the log-logistic curve – akin to a dose-response curve – provided the best fit at 21°C. The residual standard error (RSE) was smaller (5.9) in the log-logistic curve than the linear model (7.7). The inflection point of the curve was at 28.1 hours (95% CI: 24.6 – 31.5 hours), suggesting a key change at this time (Fig 3).

**Fig. 3-.**
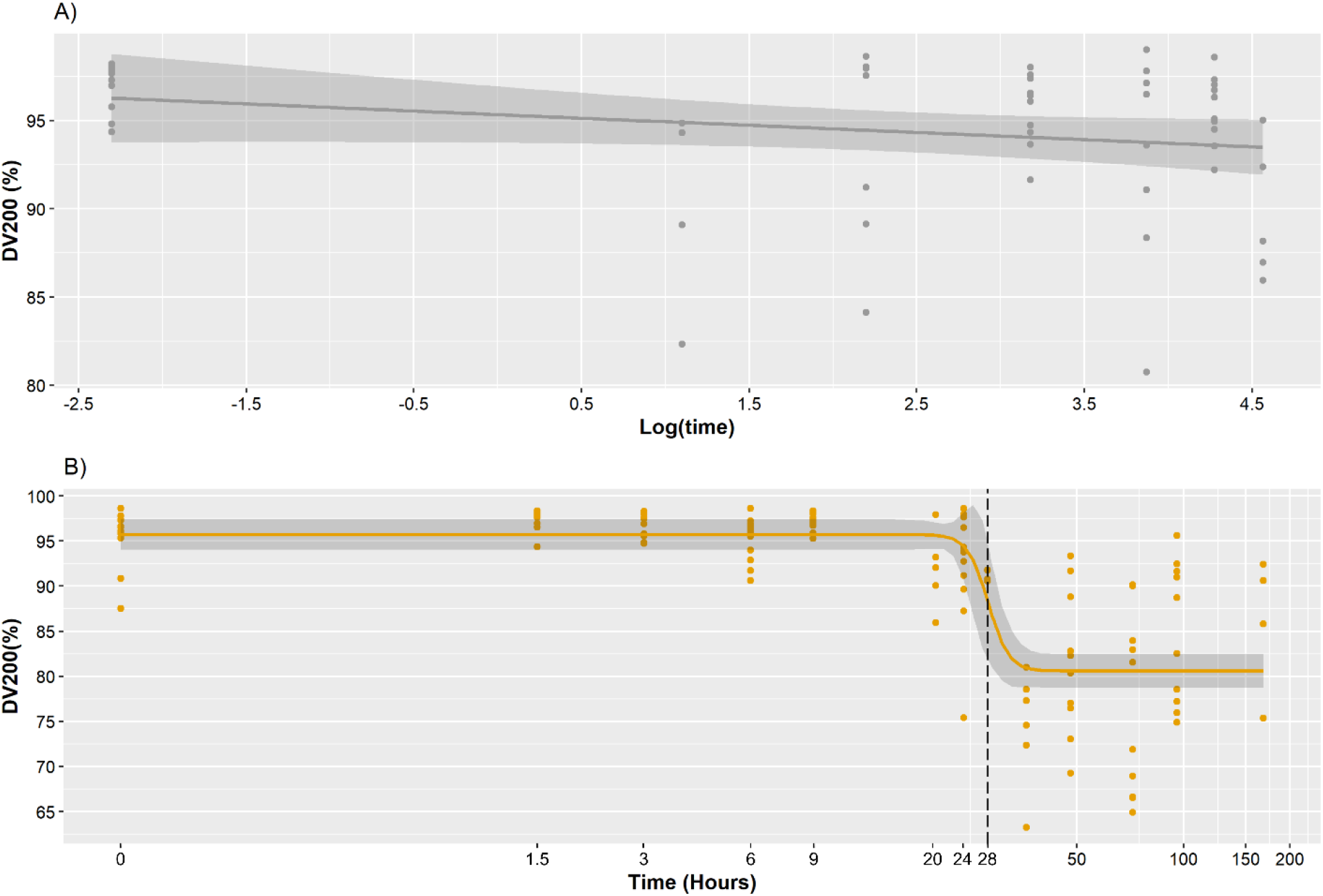
Relationship between the DV200 and time and temperature. **A)** Linear regression curve for the DV200 against log-transformed time at 4°C. A negative slope is observed. **B)** Log-logistic curve for the DV200 against time at 21°C. The curve displayed an inflection point at 28.1 hours, indicated by the vertical dashed line. Shaded areas in **A)** and **B)** represent the standard error. Note that the y-axis is scaled differently for **A)** and **B)**. The interaction effect between log-transformed time and temperature is not shown.

**Fig. 4-.**
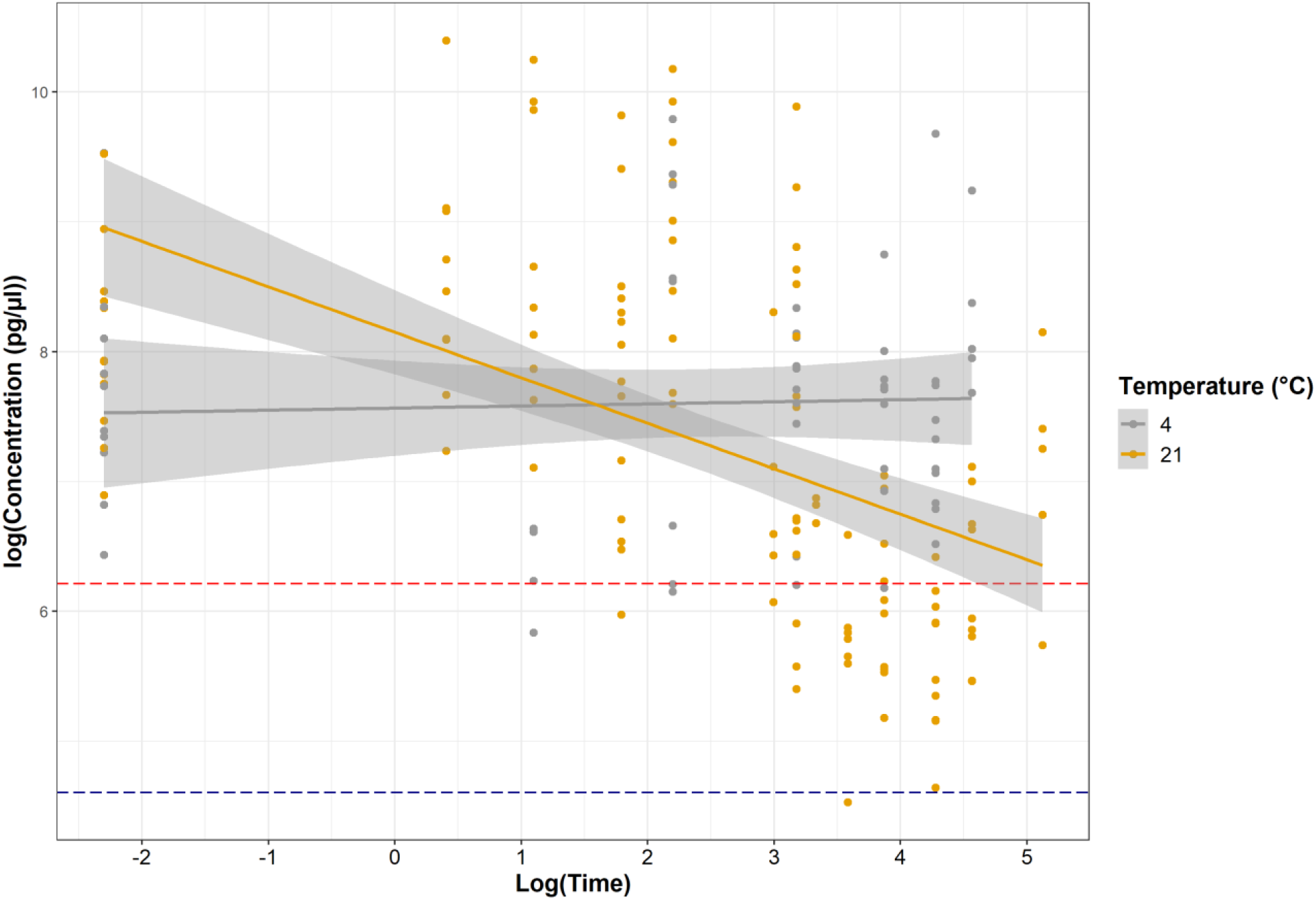
Relationships between total RNA concentration and log-transformed time (hours). Concentrations are grouped by storage temperature for comparison purposes; points and regression lines are coloured according to storage temperature. Shaded areas around the regression lines represent the standard error. Concentration was log-transformed for visualization purposes. The red dashed line represents the 4200 TapeStation’s limit of quantification (500 pg/μL) and the navy dashed line represents the instrument’s limit of detection (100 pg/μL).

### Total RNA concentration

The concentration of total RNA was shown to have a lower model fit than the RINe or DV200 (Table 2); the addition of biological replicate as a random effect greatly improved the model’s fit, suggesting high inter-individual variation (Table 2). We detected a significant relationship between total RNA concentration and time and temperature (Table 2, Fig 4). Importantly, we uncovered differences between concentrations from biological replicates originating from the same temperature (Table 3, Fig. S2). Density plots demonstrated bimodal distributions for temperature data, concordant with the result that RNA concentrations from different biological replicates at the same temperature were different (Table 3, Fig. S3). Most concentrations recorded for bloodstains deposited at early timepoints were above the limit of quantification (500 pg/μL), while many concentrations at room temperature fell below this threshold as time progressed (Fig. 4).

### 28S/18S rRNA peak area ratios

The 28S/18S rRNA peak area ratio was the retained RNA metric that displayed the weakest model fit and little relationship to time or temperature (Table 2). Peak area ratios were only obtained for timepoints up to 24 hours for bloodstains deposited at room temperature and 72 hours for those stored at 4°C (Fig. S4); the 28S/18S peak area ratios could not be obtained for the later timepoints, as the 18S rRNA peak dropped out of the electropherograms.

## Discussion

Estimating the TSD of biological fluids found at crime scenes, such as bloodstains, remains a challenge within the forensic community (1,2). Intra and inter-individual variation in blood composition, substrate type and storage conditions all influence bloodstain degradation and must be accounted for to accurately estimate TSD (2). This study provided a novel approach to the TSD of blood, by using a non-targeted, amplification-free technique within a short time since deposition period (168 hours). This is contrary to most nucleic acid studies that used targeted approaches (mRNA transcripts), relied on amplification, and explored longer time periods (3,19,20). Similarly to what Salzmann *et al.*(35) demonstrated using the integrity of mRNA transcripts (TINe – transcript integrity number equivalent), we found that total RNA quality metrics, such as the RINe and DV200, both provided clear relationships with time, with a minimal effect of biological replicate on the model. A key observation here is that the limited effect of biological replicate and inflection point at 28 hours of DV200 suggests these could be universal diagnostic metrics.

The RINe is an RNA quality metric that is equal to the ratio of the 18S rRNA peak height to the signal present in the fast zone (approximately 200 bp - 1.8 kb), which represents fragmented RNA (45). This ratio normalized the discrepancies observed in total RNA concentration and had a clear relationship to time (Fig. 2, Table 2). The addition of biological replicate as a random effect in the model only marginally increased the model fit (Table 2), which was driven by a temperature-donor effect (Table2, Table 3). This is important because previous degradation models found a large effect of biological replicate (28), but the standardized nature of RINe makes it a universally comparable metric. Blood composition varies by individual, for example, research on human blood showed that the amount of RNA in healthy subjects varied from 6.7 to 22.7 μg/ml (51). This variability could lead to significant differences in concentrations of total RNA obtained from biological replicates, as we observed during our study (Fig S2). The RINe metric was not affected by this variation (Table 2, Table 3); additionally, the RINe was not highly correlated to total RNA concentration and was independent of sample volume, highlighting the potential for wide-scale applicability. The independent and negative correlation with time suggests the utility of RINe as a useful TSD metric. We also have no reason to suspect that the patterns observed herein would not be present in human blood; bovine blood has previously been optimized for forensic research when acid dextrose anticoagulant solution A (ACD-A) is added at a concentration of 12.5% v/v, as it exhibits similar fluid properties to that of human blood (41). Bovine blood also has a similar genome size and composition to humans (52) and has similar RNA concentration (51,53), white blood cell (WBC) count (human: 3.4-11.6 × 10^3^/μL, bovine: 4.9-13.3 × 10^3^/μL), and subtypes of WBCs as human blood (neutrophils and lymphocytes are the most abundant in each organism) (51,54). Confirmatory analyses validating our results using human blood could make this technique applicable to crime scene samples in the future.

A common theme observed is the preservation effect of cold temperatures. The bloodstains at 4°C consistently demonstrated improved quality metrics when compared to the bloodstains stored at 21°C. Colder temperatures have been shown to preserve nucleic acids and slow their degradation (15,16,55), a trend that was reproduced herein by comparing the slopes observed in the linear models for the RINe and the DV200 (Fig. 2, Fig. 4A, Fig. S1). In both instances, room temperature produced steeper slopes, indicative of faster degradation. Previous studies have shown that the DV200 provides an alternative to the RINe when analyzing highly degraded samples, where electropherograms lack the 18S and 28S rRNA peaks due to fragmentation (49,56). The DV200 is often used as a quality indicator before library preparation for high-throughput sequencing, as it is not contingent on rRNA being present to provide measurements (49,57). Our results indicated that the DV200 was influenced by the dropout of 18S rRNA with time (Fig. 2); specifically, the values obtained at room temperature exhibited non-linear behaviour over time that appeared to be directly driven by the degradation of the 18S rRNA peak occurring after one full day of exposure (Fig. 2B, Fig. S5). Non-linear relationships between response variables and time have been observed in previous blood TSD studies (13–15,20), indicating that non-linearity, potentially resulting from preferential degradation of certain RNA fragments (or parts therein) (20), oxidation and dehydration (13), is not uncommon. The non-linearity and inflection point of the DV200 values at room temperature could be exploited as a diagnostic tool to differentiate bloodstains aged for less than 28 hours from older ones as there is a precipitous loss of RNA that occurs at this time (Fig. 2B), with most measurements approaching the instrument’s limit of quantification (Fig. 4), implying that little RNA is present in the sample. It is conceivable that the same curve and diagnostic signature would be present at 4°C over longer periods. Future work should extend the 4°C sampling period as this could prove useful for crime scene investigation in cold climates (58).

As this approach analyzed total RNA rather than mRNA, direct comparisons between these results and those obtained in previous studies can be difficult. Low RNA concentrations were observed throughout the study, often below the limit of quantification, as time progressed. The size bin ranging from 200 bp – 500 bp, which should represent the smaller mRNAs in the samples, followed the same trend as total RNA and produced concentrations near the limit of detection (Fig. S6). This is consistent with known half-lives of mRNA (59). Although trends in total RNA degradation tend to follow those found using mRNA transcripts, the latter is more sensitive as it targets a specific gene and can determine actual copy numbers via amplification (3,16,20). It is also susceptible to genomic DNA contamination (37). Fu *et* al. (20), and Weinbrecht *et* al. (19), both detected blood-specific transcripts from bloodstains deposited for up to one year, while the total RNA concentration in this study fell below the quantitative range after ∼24 hours of deposition. Here, we suggest the faster degradation of total RNA observed in our study allows for better discrimination at early timepoints than mRNA studies.

## Conclusion

This study demonstrated significant changes in the RINe values and a potentially diagnostic inflection point in DV200 of RNA extracted from bloodstains deposited for up to 168 hours. The biological replicates included in this study produced similar results for the quality Moving forward, in addition to an increase in sample size, longer time series at colder temperatures and experiments using human blood, the effects of humidity and substrate should be evaluated. These factors have been shown to affect the RNA degradation and drying times of bloodstains, which could cause the clear drop in RNA metrics to occur at earlier or later timepoints (16,25,60). Timewise changes in the rheological properties of blood, DNA and RNA could also be explored and combined with previously developed TSD models (61,62). This type of combinatorial research could bring the forensic community one step closer to developing a sensitive and robust TSD model for bloodstains that could one day be applied to crime scene samples.

## Acknowledgements

The authors would like to thank Reni Verasztó for helping with laboratory work and data collection. The authors would also like to acknowledge Alon Gabriel, Jesse Wolf, and Marie-Laurence Cossette of the Wildlife and Applied Genomics laboratory group at Trent University for their insights and feedback on writing, visualizations, and statistical analyses. The authors also thank Otonabee Meat Packers for providing the blood samples.

