## Supplemental table and figures for "Using total RNA quality metrics for time since deposition estimates in degrading bloodstains" (from another version of this preprint: version 3, 2022-04-18)

### Supplementary Materials

**Table S1.** Two-sample t-tests between RINe, DV200 and concentration of total RNA obtained from blood containing anticoagulant (ACD-A) and blood without it at each timepoint examined. 0 hours indicates the time of deposition. Blood collected from the same cow underwent one of two anticoagulant treatments: a) 12.5% ACD-A or b) no anticoagulant. 50 µL of blood was immediately deposited into 1.5 mL plastic microcentrifuge tubes upon arrival to the laboratory. All samples were left to air dry at room temperature (~21°C) by keeping the lids on the microcentrifuge tubes open until RNA extraction. RNA extractions were completed at 0 hours (time of initial deposition), 24 and 168 hours after deposition. For both treated and untreated blood, five replicates were used at each timepoint, resulting in a total of 15 samples for the treated and untreated blood, respectively. RNA extracts were analyzed following the protocol outlined in the Materials and Methods.

| RNA Metric | Timepoint | Test Statistic (t) | p-value |
|---|---|---|---|
| **RINe** | | | |
| | 0 hours (time of deposition) | -1.45 | 0.191 |
| with ACD-A vs. without ACD-A | 24 hours | -0.23 | 0.826 |
| | 168 hours | -1.27 | 0.245 |
| **DV200 (%)** | | | |
| | 0 hours (time of deposition) | 1.69 | 0.185 |
| with ACD-A vs. without ACD-A | 24 hours | 0.33 | 0.752 |
| | 168 hours | -1.61 | 0.161 |
| **Concentration (pg/µL)** | | | |
| | 0 hours (time of deposition) | 1.96 | 0.093 |
| with ACD-A vs. without ACD-A | 24 hours | 1.41 | 0.223 |
| | 168 hours | 5.34 | **<0.001** |

**Table S2.** RNA metrics provided by the 4200 Agilent TapeStation. Bolded metrics represent those that were manually added during our experiments.

| TapeStation RNA metrics | Assessment |
|---|---|
| RNA Integrity Number equivalent (RINe) | Quality |
| DV200 | Quality |
| 28S/18S rRNA peak area ratio | Quality |
| Total RNA concentration | Quantity (pg/µL) |
| 28S rRNA peak concentration | Quantity (pg/µL) |
| 18S rRNA peak concentration | Quantity (pg/µL) |
| 28S rRNA peak fluorescent units | Quantity (FU) |
| **mRNA concentration (200 – 500 bp)** | Quantity (pg/µL) |
| **mRNA fluorescent units** | Quantity (FU) |

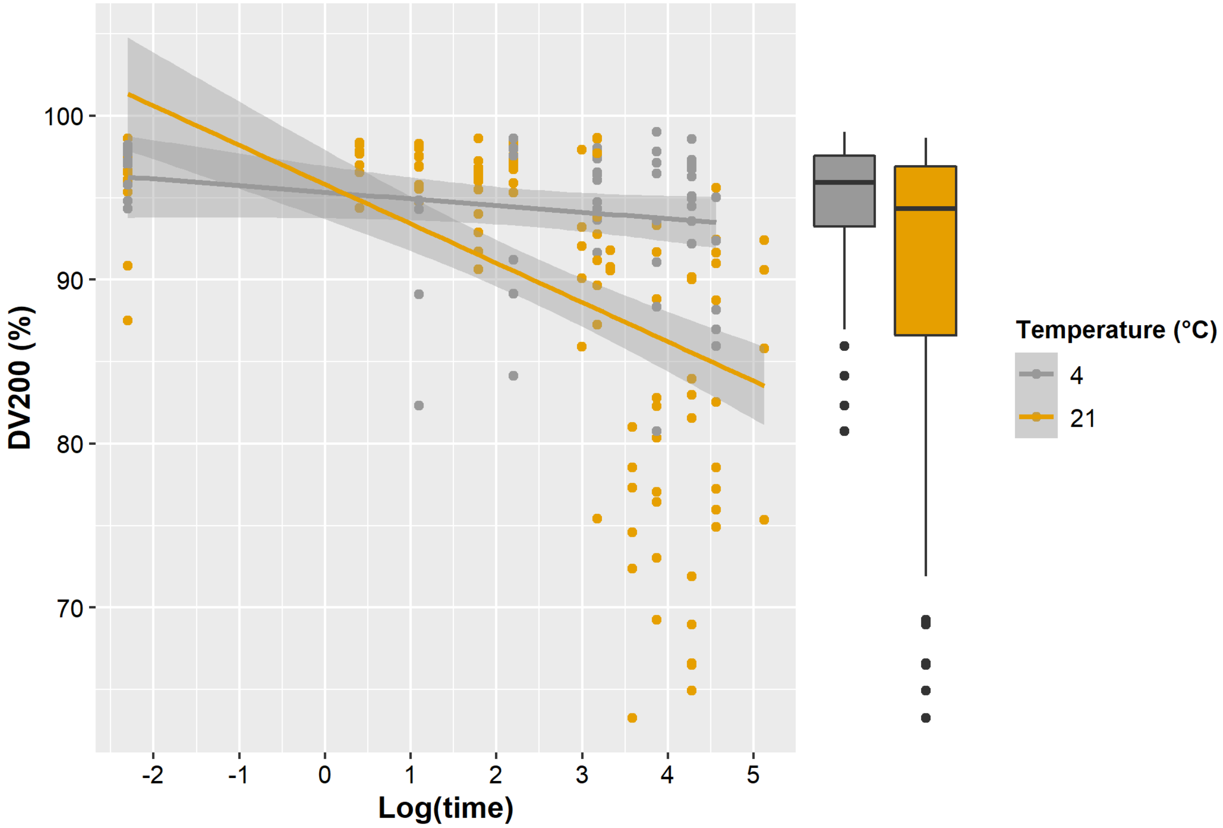


**Fig. S1.** Relationship between the percentage of RNA fragments larger than 200 bp (DV200), and log-transformed time. Points and regression lines are coloured according to the temperature at which the bloodstains were stored. Shaded areas around the regression lines represent the standard error.

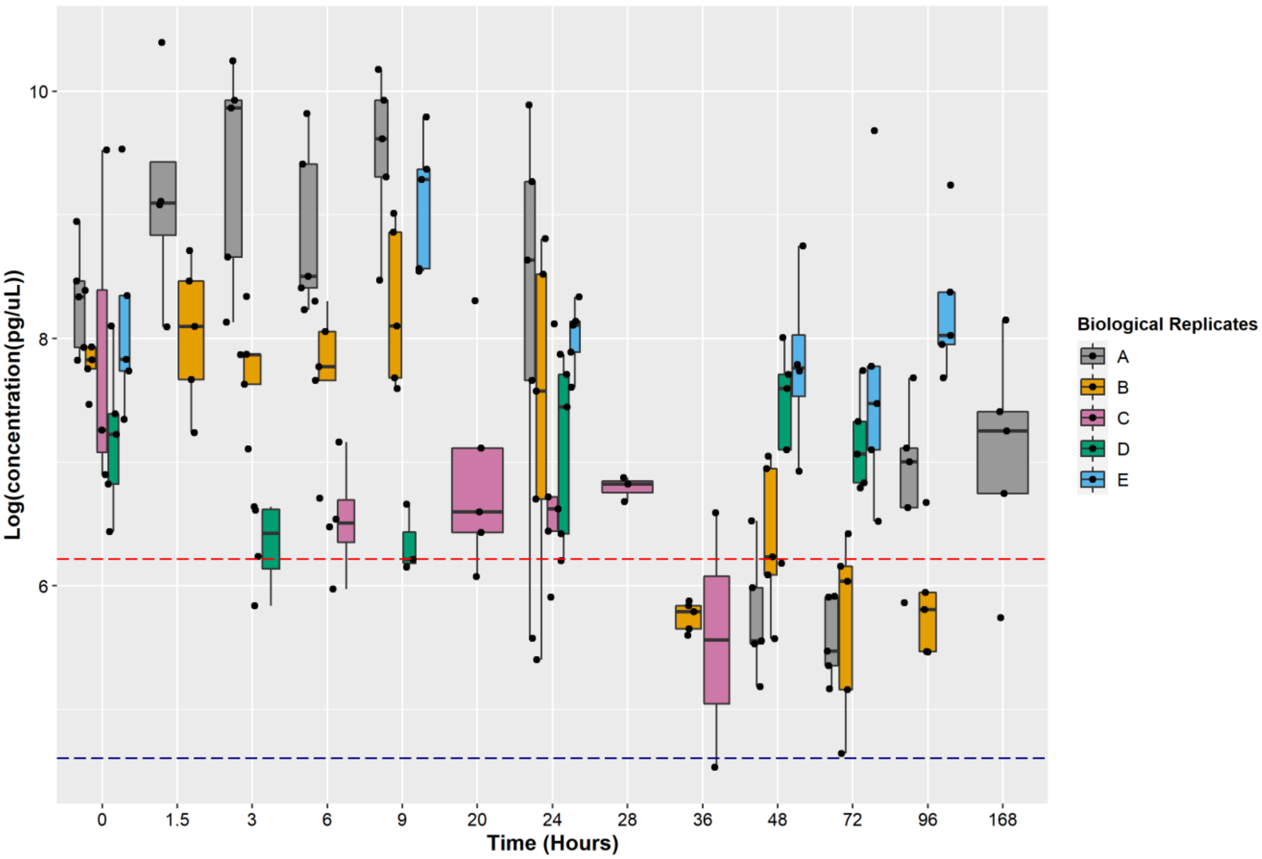


**Fig. S2.** Relationship between total RNA concentration in time. Biological replicates A-C represent the three 21˚C time series, while biological replicate D and E represent the two time series completed at 4˚C. The red dashed line represents the 4200 TapeStation's limit of quantification (500 pg/µL) and the navy dashed line represents the instrument's limit of detection (100 pg/µL).

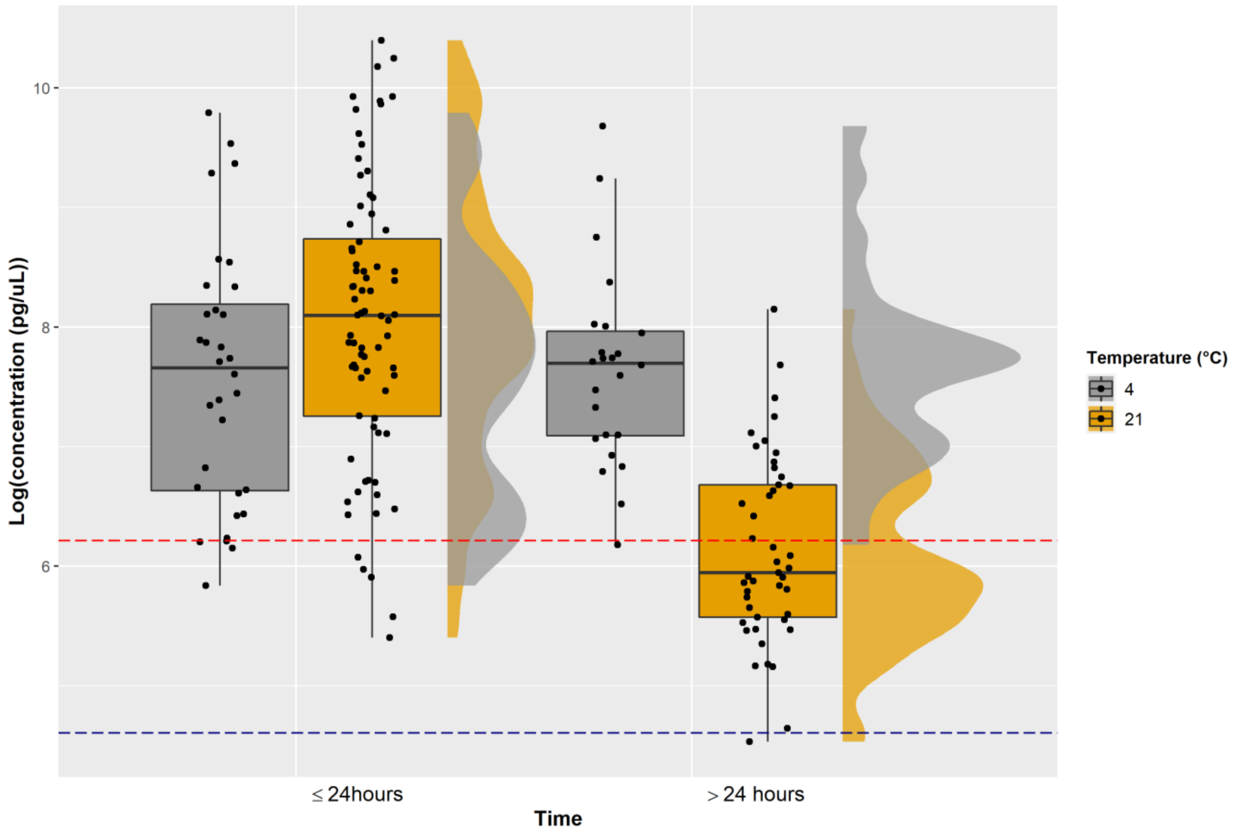


**Fig. S3 -** Boxplots and density plots illustrating the distribution of total RNA concentration (natural log-transformed) for bloodstains deposited for 24 hours or less and those deposited for longer periods of time. Raw concentration data are overlaid on the boxplots. The red dashed line represents the 4200 TapeStation's limit of quantification (500 pg/µL) and the navy dashed line represents the instrument's limit of detection (100 pg/µL).

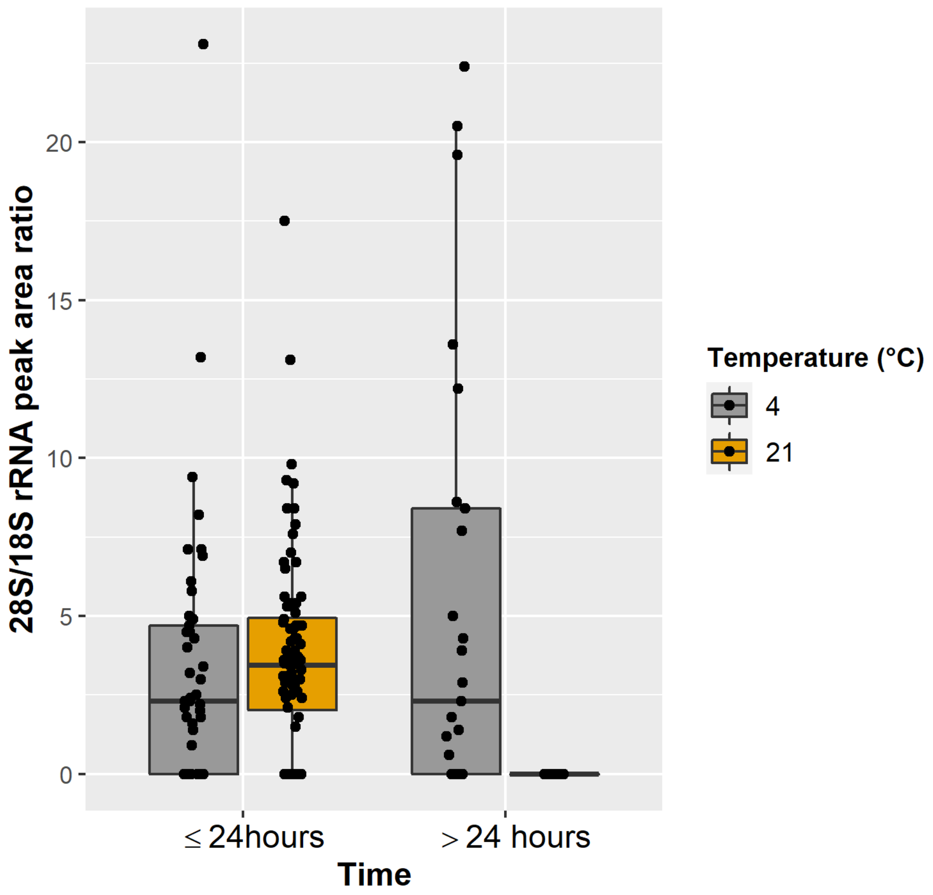


**Fig. S4.** Relationship between 28S/18S rRNA peak area ratios and time. Dropout of 18S rRNA after 24 hours for room temperature samples resulted in the inability to obtain ratios for the timepoints greater than 24 hours. Samples stored at 4 ˚C produced ratios past 24 hours.

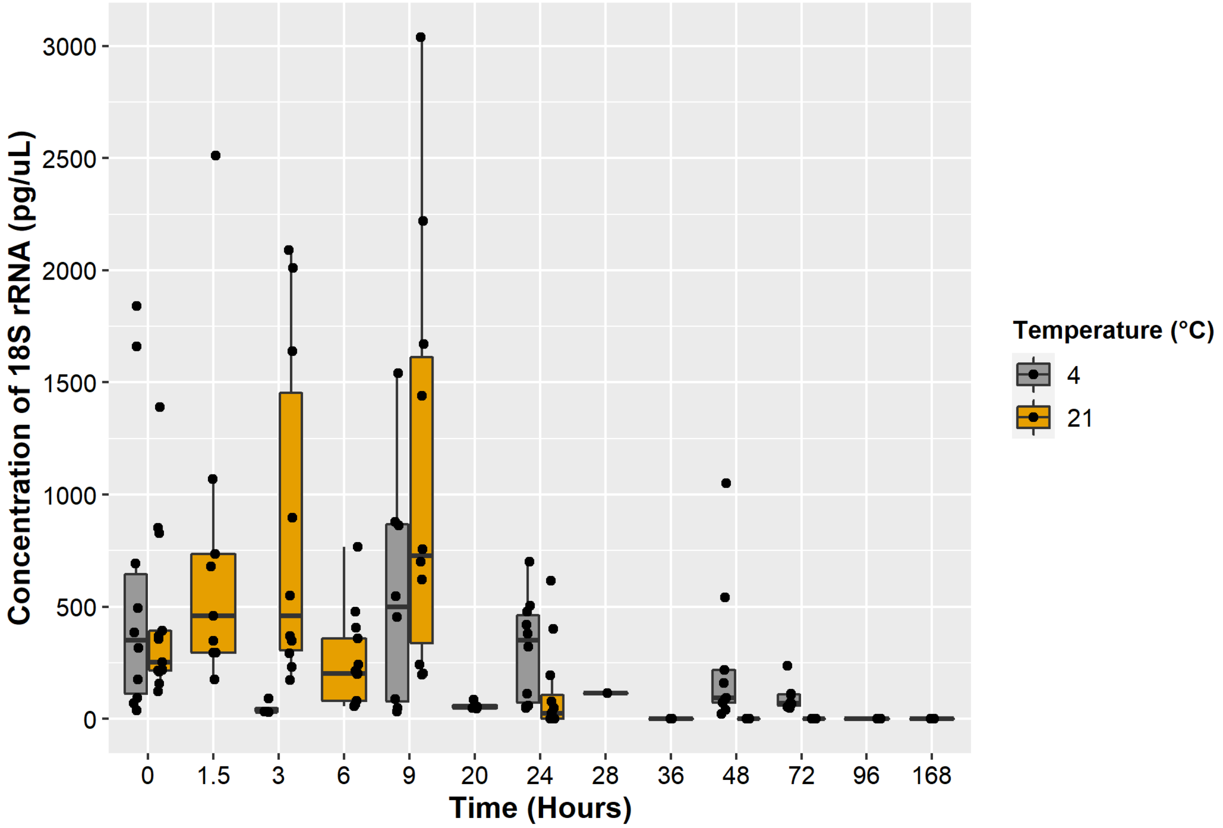


**Fig. S5.** Relationship between the concentration of 18S rRNA and time. Boxplots were coloured according to the bloodstain's storage temperature.

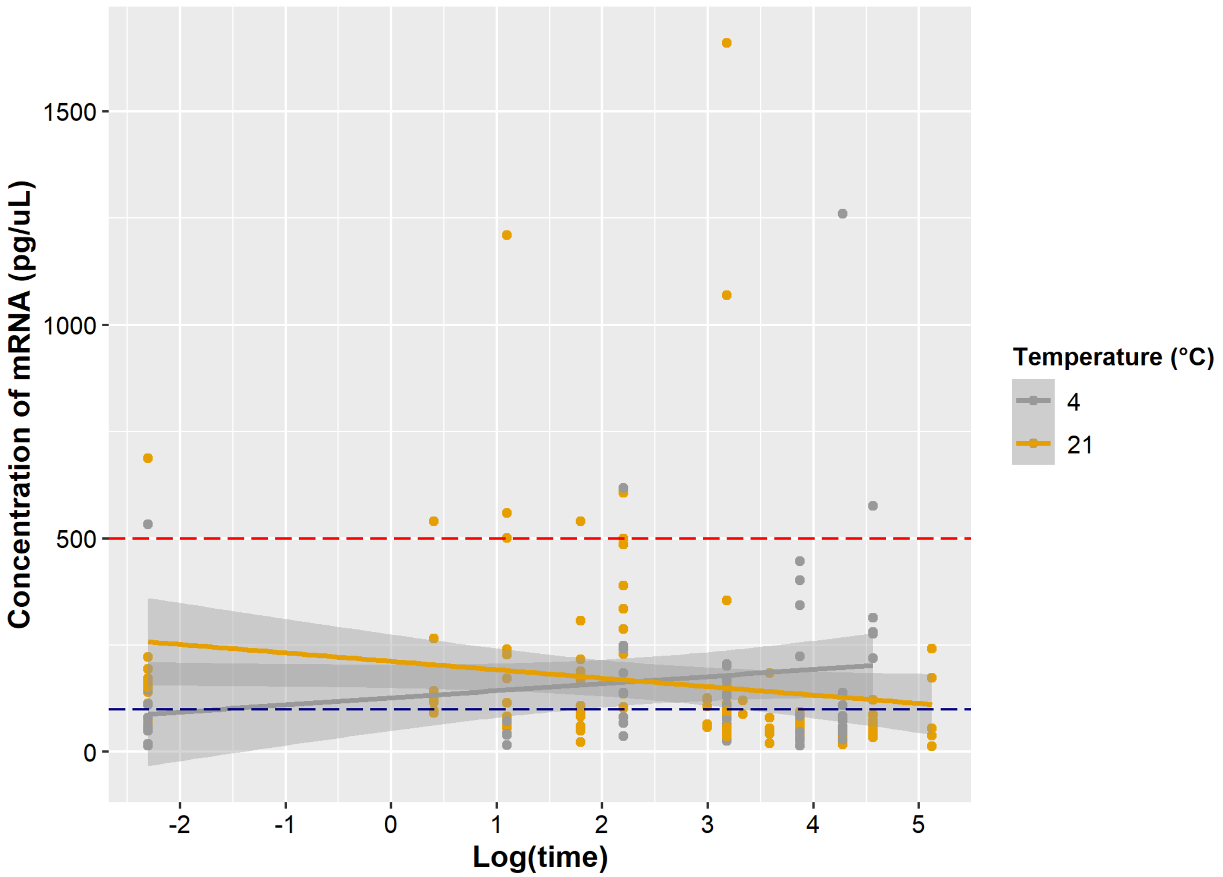


**Fig. S6.** Relationship between the 200 bp – 500 bp size bin, representing the smaller mRNAs in our sample, and log-transformed time. The red dashed line represents the 4200 TapeStation's limit of quantification (500 pg/µL) and the navy dashed line represents the instrument's limit of detection (100 pg/µL).
